# A mathematical model of heat shock response to study the competition between protein folding and aggregation

**DOI:** 10.1101/2020.04.13.039123

**Authors:** Sushmita Pal, Rati Sharma

**Affiliations:** Department of Chemistry, Indian Institute of Science Education and Research (IISER) Bhopal Bhopal Bypass Road, Bhauri, Bhopal 462066 INDIA

## Abstract

Proteins, under conditions of cellular stress, typically tend to unfold and form lethal aggregates leading to neurological diseases like Parkinson’s and Alzheimer’s. A clear understanding of the conditions that favor dis-aggregation and restore the cell to its healthy state after they have been stressed is therefore important in dealing with these diseases. The heat shock response (HSR) mechanism is a signaling network that deals with these undue protein aggregates and aids in the maintenance of homeostasis within a cell. This framework, on its own, is a mathematically well studied mechanism. However, not much is known about how the various intermediate mis-folded protein states of the aggregation process interact with some of the key components of the HSR pathway such as the Heat Shock Protein (HSP), the Heat Shock transcription Factor (HSF) and the HSP-HSF complex. In this article, using kinetic parameters from the literature, we propose and analyze two mathematical models for HSR that also include explicit reactions for the formation of protein aggregates. Deterministic analysis and stochastic simulations of these models show that the folded proteins and the misfolded aggregates exhibit bistability in a certain region of the parameter space. Further, the models also highlight the role of HSF and the HSF-HSP complex in reducing the time lag of response to stress and in re-folding all the mis-folded proteins back to their native state. These models therefore call attention to the significance of studying related pathways such as the HSR and the protein aggregation and re-folding process in conjunction with each other.

## I. INTRODUCTION

Survival under stressful conditions is one of the primary aspects of a thriving species. In general, cells face various kinds of stresses such as Heat [18], Oxidative stress [39, 43], Endoplasmic Reticulum stress [11] or stresses induced by chemical agents[10]. Therefore, living organisms have evolved various defense mechanisms to respond to such stresses. Heat stress, in particular, leads to protein unfolding and misfolding, eventually causing protein aggregation in the cell. Such aggregation impairs protein function for normal cellular processes [23, 29]. Response to this therefore requires re-folding these proteins to their native state, which is achieved through a mechanism known as the heat shock stress response (HSR) pathway.

In broad terms, the HSR pathway follows the titration feedback mechanism [1]. Under unstressed conditions, Heat Shock Proteins (HSPs), which are small molecular chaperones, exist in the form of a complex with Heat Shock Factors (HSFs). On accumulation of misfolded/unfolded proteins in the cell, the HSPs are titrated away from the HSF-HSP complex and bind to the hydrophobic residues of the unfolded proteins to fold them back. The free HSFs now dimerize, trimerize, get phosphorylated (activated) and through the process of transcription and translation produce more HSPs. These HSPs can now fold back more of the accumulated protein aggregates.

Modeling of HSR has closely followed experimental evidences that revealed pieces of the mechanism right from the beginning. The modeling frameworks have explained data from heat shock experiments in various species such as bacteria [4, 13, 16, 40], yeast [15, 46], worms [3], plants [22, 36, 37] and mammalian cells [27, 28, 30, 35, 38]. The earliest HSR model that fit the data from heat shock experiments in mammalian cells [27], introduced a mathematical framework of the titration feedback loop. Following this, there were several other models which included varying degrees of details into them, such as, transcriptional regulation [30], translation and activation of HSF [28, 35] and later a more simplified model [38].

Although the heat shock experiments have revealed the HSR mechanism, they say nothing about the different mis-folded states of the proteins themselves. In fact, many proteins are known to undergo spontaneous aggregation. This happens during amyloid formation that begins with a nucleation phase and then goes through a series of intermediate elongation steps and self propagates to spontaneously aggregate [12]. Such protein aggregation has been seen, for example, in *β*-lactoglobulin and its peptides when they spontaneously aggregate to form amyloid fibrils [9]. Spontaneous aggregation also occurs in PHF6, a key hexapeptide of tau protein, the mechanism for which was shown in a recent molecular dynamics simulation study [21]. In addition to such aggregation, proteins are also known to show hysterisis between unfolding and folding. This kind of bistability (the presence of two stable states of proteins) is exhibited in the guanidine hydrochloride induced unfolded and refolded states of Transthyretin [17] and MTase proteins [2]. Such threshold crossing is therefore quite common in protein folding and aggregation.

The above mentioned mathematical models of the HSR explained the various mechanistic details. However, these did not look into the the folded and aggregated states of the proteins themselves. In recent years, there have been a few simplified models which looked at the interplay between folded and aggregated states [26, 31, 34]. These models represent over-simplified threshold cross-over (or bistability) between folded and aggregated states. The first of these three models only included the HSP counts and ignored every other feature of the HSR framework, while the latter models did not even include that. Therefore, it is evident that more work needs to be carried out to explore the bistable phenomenon in conjunction with HSR, where the interaction network is not just restricted to the folded and aggregated states, but also includes specific entities of the HSR pathway, such as, HSF, HSP and the HSF-HSP complex.

In light of this, we have in this work, explored bistability in the competition between folded and aggregated states while still incorporating features of the HSR framework. We included interaction of proteins not just with HSP but also with the HSF-HSP complex in addition to activation of HSF in response to heat shock. We explore these features through two specific models with increasing levels of detail.

Our main findings from these explorations are that the presence of HSF and HSF-HSP complex is important for conversion of misfolded proteins to theirnative folded state. Further, there is delayed response to heat shock which manifests as a slowly decreasing aggregated steady state which then crosses the threshold of bistability to become completely folded. This feature only results when there is production of more HSP through HSF activation due to increased stress and not otherwise. We develop the mathematical models through a system of ODEs and then carry out stochastic simulations to explore the probability distributions of the folded and aggregated states of the protein. The methods used are discussed in Section II. Section III gives a detailed account of the specific results obtained via the two models, while Section IV provides discussions and implications regarding the larger context of these results.

## II. MATHEMATICAL MODEL AND METHODOLOGY

### Model Structure

The basic pathway of the heat shock response is as follows. The cell activates its response mechanism when it receives a stress signal. In the case of heat stress, the response is elucidated as increased production of heat shock proteins (HSPs) [6], which are small molecular chaperones that act to fold back mis-folded proteins. The HSPs usually exist in the form of a complex with the Heat Shock Transcription Factor (HSF). Unfolded proteins titrate away the HSPs from this complex. Increased stress causes more proteins to become unfolded thereby increasing the demand for HSPs. This increased production of HSPs is facilitated through activation of free HSFs. The mis-folded proteins also go through several intermediates, both during the aggregation and the re-folding processes. In this study, we develop models that capture these intermediate states along with the HSR mechanism.

Taking into consideration the folding and aggregation process of proteins along with the HSR pathway, we propose and analyze two possible models for protein aggregation in this study. Both these models incorporate HSF and its interaction with HSP along with the different stages of the folding pathway, something that had not been considered before. In Model I, we assume that the total HSPs and HSFs in the system are conserved. Model II is an extension of Model I where we let the HSPs be dynamically produced. We consider two models because they represent varying degrees of detail in the HSR pathway, as will become clear when we explore the individual models through simulations. These models give a better understanding of the Heat Shock Response and and elucidate how HSPs are produced when the cell is exposed to Heat Shock. A combined picture of Models I and II is shown in Figure 1 and the reactions specific to individual models are listed in Table I.

**FIG. 1:**
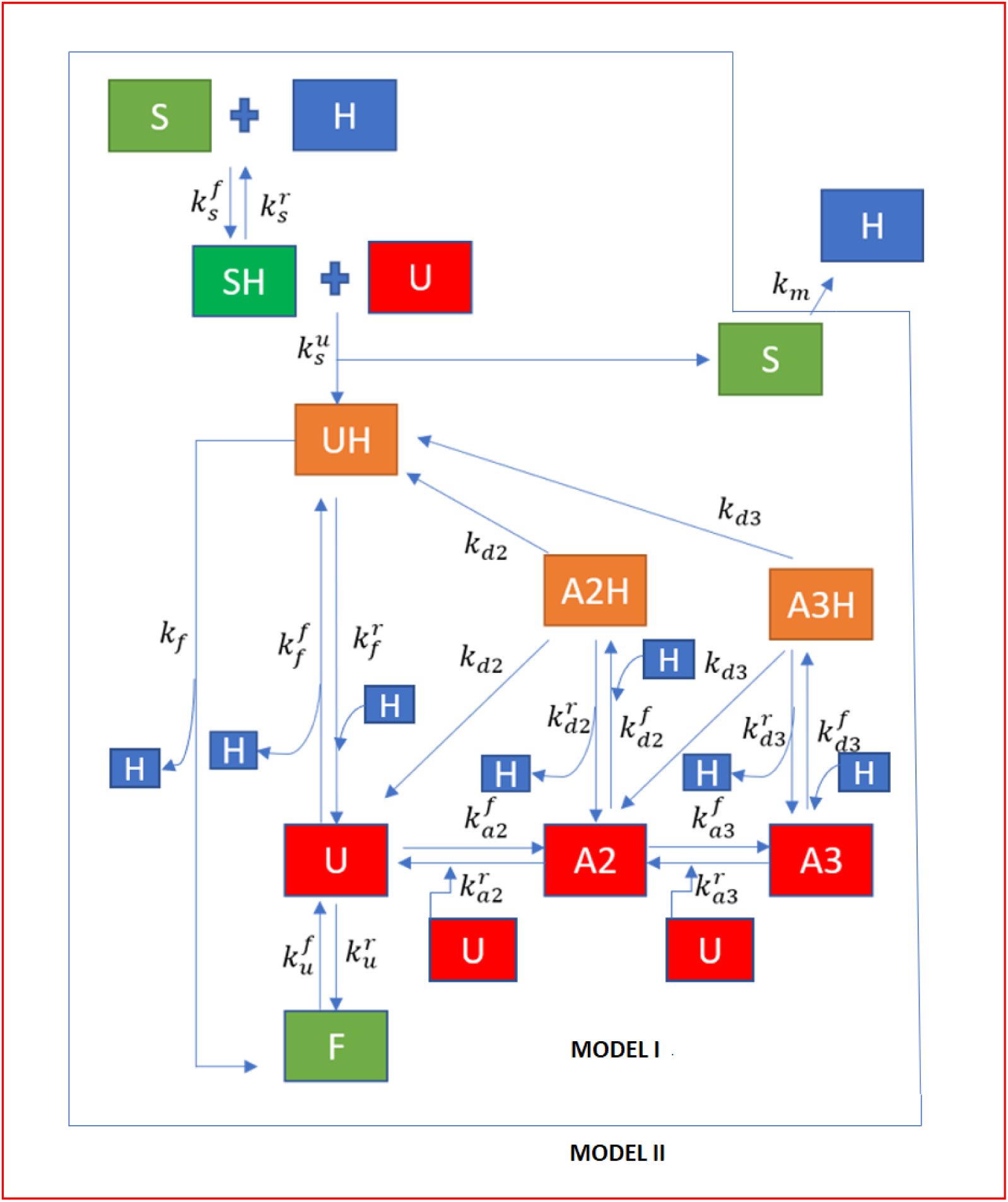
Our mathematical model. The single headed arrows represents irreversible reaction whereas single headed two arrows running opposite to each other represent reversible reactions. U, A2, A3 are unfolded, aggregate with 2 monomers and aggregate with 3 monomers respectively. UH, A2H, A3H are unfolded proteins and aggregates bound to HSP. S, H, F, SH are HSF, HSP, Folded Protein and HSF:HSP complex respectively.

**TABLE I:** The complete list of reactions corresponding to each model.

|  | Model I | Model II |
| --- | --- | --- |
| **Reactions** | $F \rightleftharpoons U$ $U + U \rightleftharpoons A_2$ $A_2 + U \rightleftharpoons A_3$ $U + H \rightleftharpoons U : H$ $A_2 + H \rightleftharpoons A_2:H$ $A_3 + H \rightleftharpoons A_3:H$ $A_2:H \implies U + U : H$ $A_3:H \implies A_2 + U:H$ $U:H \implies F + H$ $S:H \rightleftharpoons S + H$ $S:H + U \implies U : H + S$ | $F \rightleftharpoons U$ $U + U \rightleftharpoons A_2$ $A_2 + U \rightleftharpoons A_3$ $U + H \rightleftharpoons U : H$ $A_2 + H \rightleftharpoons A_2:H$ $A_3 + H \rightleftharpoons A_3:H$ $A_2:H \implies U + U : H$ $A_3:H \implies A_2 + U:H$ $U:H \implies F + H$ $S:H \rightleftharpoons S + H$ $S:H + U \implies U : H + S$ $S \implies H$ |

The models proposed consider aggregate formation from upto 3 unfolded monomers of protein, taking into consideration that the higher degrees of aggregation show a similar pattern of aggregation. Such a model helps us examine the unique behaviour of two distinct processes i.e, monomer-monomer dimerization and monomer-oligomer dimerization and the effect of their rate constants on bistability. In a real cell, HSPs exist in several different forms such as HSP16, HSP70, HSP90, etc [5, 6]. Similarly, many different kinds of proteins can be folded back by these chaperones when they get mis-folded. However, for the sake of simplicity and as per standard practice, we choose not to differentiate between various kinds of HSPs and proteins in both these models. Therefore, in our models, HSPs, folded proteins (F) and unfolded proteins (U) serve as stand-ins for several possible different molecules of the same class.

### Parameter estimation

As listed in Table II, we chose our parameters from literature on mammalian cells. We found that Robinson et al’s [33] work on Chaperone activity and protein folding process inside eukaryotic cells provided us with the range of values for several of our parameters. In order to study the catalytic role of chaperone in protein folding, we also assumed self re-folding of unfolded proteins to be several orders of magnitude lower than chaperone assisted refolding [31].

**TABLE II:** List of parameters. The values of rate has units of (*item × s*)^−1^ for a second order reaction and *s*^−1^ for a first order reaction.

| Parameter | Value [<br>(*item* × *s*)<sup>-1</sup><br>or *s*<sup>-1</sup> ] | Description | Reference |
| --- | --- | --- | --- |
| $k_u^f$ | 10<sup>-4</sup> | Temperature dependent misfolding of folded proteins | Eq. 1 |
| $k_u^r$ | 10<sup>-5</sup> | Self Refolding of misfolded proteins | Robinson et al,1996 [33] |
| $k_{a2}^f$ | 0.02 | Rate of aggregation of two U | Zettlmeissel et al,1979 [45] |
| $k_{a2}^r$ | 0.04 | Self disaggregation of $A_2$ | Rieger et al,2006 [31] |
| $k_{a3}^f$ | 0.01 | Rate of aggregation of $A_2$ and U | Zettlmeissel et al,1979 [45] |
| $k_{a3}^r$ | 0.01 | Self disaggregation of aggregate $A_3$ | Rieger et al,2006 [31] |
| $k_f^f$ | 0.01 | Rate of U and H binding | Flynn et al.,1989 [7] |
| $k_f^r$ | 0.1 | Rate of disaggregation of U:H | Flynn et al., 1989 [7];<br>Liberek et al., 1991 [20] |
| $k_{d2}^f$ | 0.1 | Rate of entrapment of H with $A_2$ | Robinson et al,1996 [33] |
| $k_{d2}^r$ | 0.1 | Rate of disaggregation of $A_2$:H | Robinson et al,1996 [33] |
| $k_{d3}^f$ | 0.001 | Rate of entrapment of H with $A_3$ | Robinson et al,1996[33] |
| $k_{d3}^r$ | 0.1 | Rate of disaggregation of $A_3$:H | Robinson et al,1996 [33] |
| $k_{d2}$ | 3×10<sup>-3</sup> | Rate constant for degradation of $A_2H$ to U and U:H | Rieger et al,2006 [31] |
| $k_{d3}$ | 3×10<sup>-3</sup> | Rate constant for degradation of $A_3H$ to $A_2$ and U:H | Rieger et al,2006 [31] |
| $k_f$ | 10<sup>-3</sup> | Catalytic refolding of proteins from UH | Rieger et al,2006 [31] |
| $k_s^r$ | 3.56 | Rate of dissociation of S:H into S and H | Petre et al, 2011 [28] |
| $k_s^f$ | 9.73 | Rate of association of S and H | Petre et al, 2011 [28] |
| $k_s^u$ | 10<sup>-4</sup> | Rate of S:H binding to U | Mikolaj et al, 2013 [41];<br>Zuzanna et al, 2009 [47] |
| $k_m$ | 10<sup>-4</sup> – 10<sup>-3</sup> | Rate of HSP translation | Scheff et al, 2015 [35], Petre et al 2011 [28],<br>Sivory et al 2016 [38] |

**TABLE III:** Sensitivity coefficients describing the role of parameters in the counts for various species. The listed values are the ones that significantly alters the processes inside the cell and are most sensitive.

| Description | Sensitivity Coefficient | Value |
| --- | --- | --- |
| Temperature induced mis-folding of folded proteins | $\frac{\partial \ln f}{\partial \ln k_u^f}$ | -1 |
| Forward aggregation decreases the complex U:H | $\frac{\partial \ln uh}{\partial \ln k_{a3}^f}$ | -0.550 |
| Smaller aggregates tend to decrease as chaperones bind to higher aggregates | $\frac{\partial \ln a_2}{\partial \ln k_{d3}^f}$ | -0.557 |
| Folded proteins increases as chaperones bind to the smaller aggregates | $\frac{\partial \ln f}{\partial \ln k_{d2}^f}$ | 0.506 |
| Chaperones assisted refolding enhances its binding to aggregates | $\frac{\partial \ln a_{2h}}{\partial \ln k_f^f}$ | 0.769 |
| Folded proteins increases as S:H binds to unfolded proteins | $\frac{\partial \ln f}{\partial \ln k_s^u}$ | 0.158 |
| Folded proteins increases as the rate of S and H binding increases | $\frac{\partial \ln f}{\partial \ln k_s^f}$ | 1.18 |
| U:H complex increases as the rate of S and H binding increases | $\frac{\partial \ln uh}{\partial \ln k_s^f}$ | 1.18 |
| S:H complex increases as the rate of S and H binding increases | $\frac{\partial \ln sh}{\partial \ln k_s^f}$ | 7.46 |

The stress was introduced in the model by specifying the value of the parameter 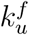, i.e., the rate at which the folded proteins unfold. Previous studies performed on various mammalian cells [19, 27], show the nature of denaturation of proteins as a function of time. The proteins in these experiments were exposed to gradually increasing temperature and the differential scanning calorimetric data was procured for transition or melting temperatures. The curve obtained from the fractional protein denaturation as a function of temperature was approximated to obtain Eq. 1. The parameter 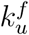, according to this equation, increases as a power law with increase in temperature. This equation depicts the rate of denaturation in the temperature range of 37-45 °C.

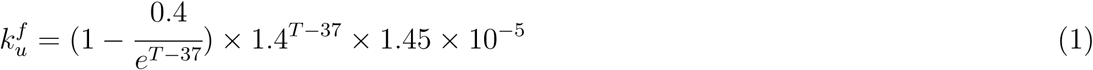

In experimental studies of the HSR mechanism, cells are made to undergo hyperthermia, which is defined as the exposure of cells to temperatures above 37°C. In line with these experimental studies, hypothetical heat stress induced in our model was around 40°C and Eq. 1 was used to calculate 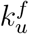 at this temperature. Following the above basic principles, we compiled the complete set of parameters. These are listed along with their respective references in Table II.

### Deterministic analysis and stochastic simulations

As a first step to defining our models, we wrote down the chemical kinetic rate equations corresponding to the changes in flux of the individual components undergoing the reactions. This system of ordinary differential equations (ODEs) is shown in Equations 2-11. As mentioned in Table II, the kinetic rate parameters were chosen from literature. The descriptions of the various kinetic parameters of the models are also stated in Table II. We then solved this system of equations at steady state and determined whether the folded, unfolded and aggregated proteins exhibited bistability. We further explored this bistability through stochastic simulations.

Exploring the system through simulations is important because reactions taking place inside a living cell are associated with stochasticity due to both population heterogeneity of the various components of the reaction system and the fluctuations inherent in the cellular environment. This inherent stochasticity in the system leads to fluctuations both in the reaction time and in the copy number of individual species undergoing the reactions. Therefore, a need for carrying out stochastic simulations arises in order to account for these fluctuations. The magnitude of these fluctuations (also called noise), has an inverse relationship with system size. This is reflected in the copy number fluctuations in our system.

The system under consideration exhibits bistability. This means that the system fluctuates between two stable states through an unstable state. The switch between the two states occurs when a fluctuation around one stable state pushes the system across the threshold taking it to the other stable state. This switch can be seen in the individual trajectories of the simulations (Figs. 3 through 8).

Therefore, it is of utmost importance to perform stochastic simulations to get a better insight into the bistable behaviour of the system. The stochastic behaviour can be studied by writing down the corresponding rate of change of probabilities of the system being in a certain composition with respect to its components. This equation, known as the Chemical Master Equation (CME), cannot be solved analytically for a system of this level of complexity. Stochastic simulations of such systems using the Gillespie algorithm [8] therefore provides an alternative.

The simulations in this study were run for 1000 replicates to study the noise in the system. The resulting probability distribution of the copy numbers of all the components were plotted to see the effect of chaperones and HSF dynamics in determining the state of the cell. The simulations themselves were performed using the simulation software package, Lattice Microbes [32] which incorporates the Gillespie algorithm. The simulations for both the models were carried out by setting up every reversible reaction as two individual independent reactions. The stoichiometric matrix and dependency matrix were explicitly written to describe the nature of the reactions and the effect on species count after every time step.

## III. RESULTS

We have, in this article, presented two models (I and II), which are representations of the protein folding and aggregation pathway with increasing levels of details from the HSR pathway. We analyze the two models to show how incorporating features from the HSR pathway makes for a better model for the interaction of the HSPs with the unfolded proteins.

### A. Model I

Model I is represented in Table I and its mathematical equations representing the change in concentrations of the individual species are represented in Equations (2) through (11).

#### Deterministic analysis

We first carried out a deterministic analysis of Model 1. The conservation of total proteins, total HSP and total HSF are represented mathematically in Equations (2) to (4). The kinetic rate equations of the individual species are shown in Equations (5) through (11).

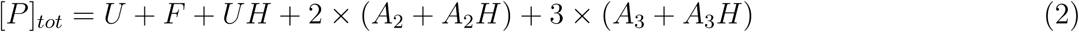

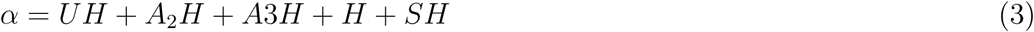

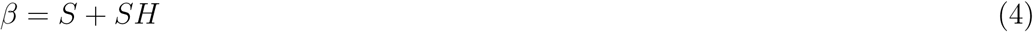

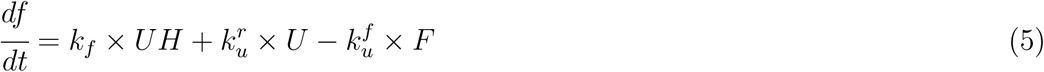

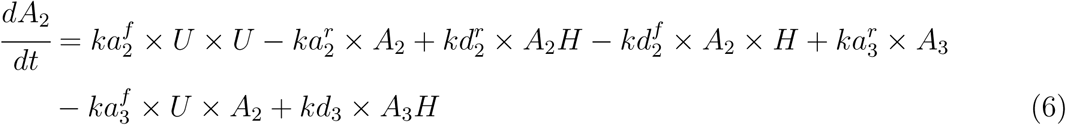

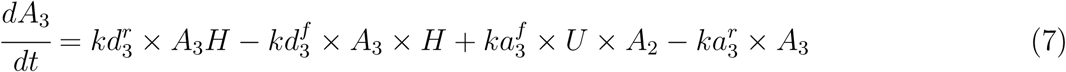

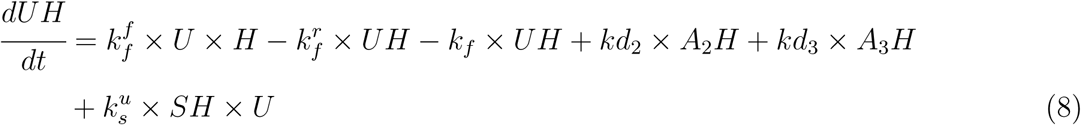

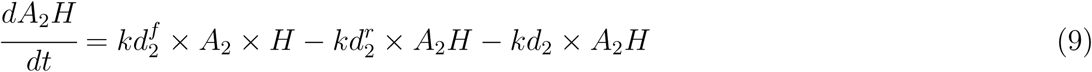

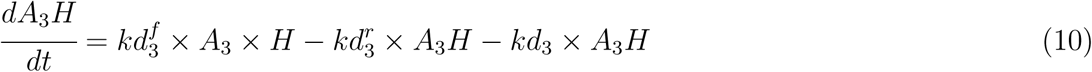

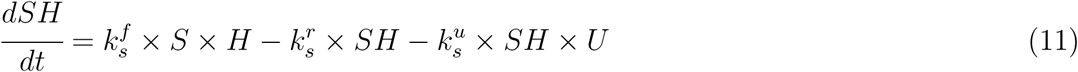

We solved the system of ODEs along with the constraints, i.e., Eqs. (2) through (11), under steady state conditions (where the change in copy numbers with respect to time is set to zero) to find regions of parameter space that have 3 or 1 physically relevant solutions for the copy numbers of folded (*F*), unfolded (*U*) and aggregated (*A*_2_ and *A*_3_) proteins. The region of space that has 3 steady state solutions shows bistable behavior while that with only one steady state solution shows monostable behavior.

Since the total HSP and HSF are conserved in Model I, the components of the system in the bistable regime can potentially redistribute itself in such a way that there are two stable steady states with an unstable state in the middle. The deterministic analysis was explicitly performed for Model 1. The corresponding parameter scan with the two regions (bistable and monostable) explicitly displayed are plotted in Fig 2. Following this, stochastic simulations were also carried out.

**FIG. 2:**
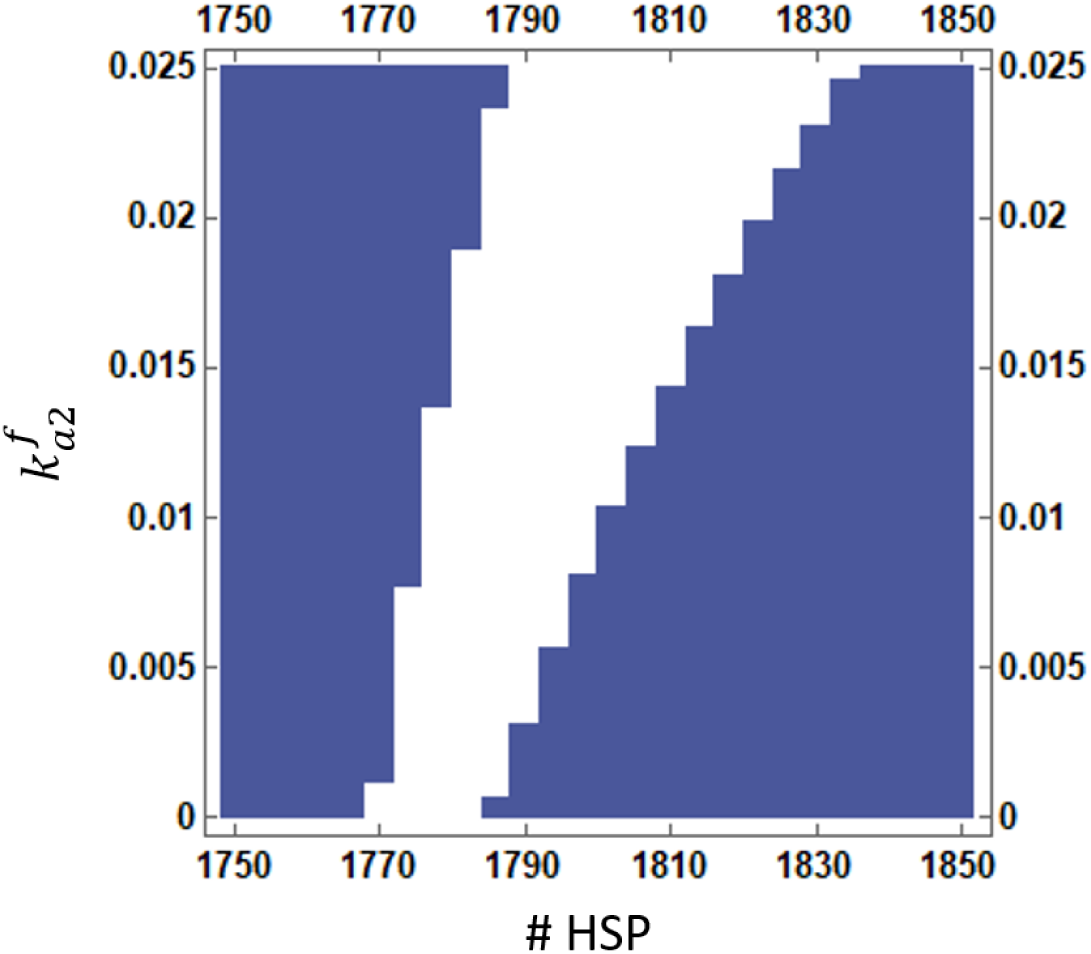
Parameter scan to determine the bistable regime. The white shaded region of the figure shows the bistable regime where the system has of 3 steady states. The system of ODEs were solved with a two parameter scan over a range in parameter space. 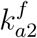 and concentration of HSP were the two parameter chosen for scan. The total protein count, S or SH count is 5000 and 1000 respectively.

#### Bistability

We varied the total HSP and the kinetic rate constant for aggregate formation, 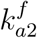, to carry out the two parameter scan in order to determine the number of physically relevant solutions of the ODEs at steady state. We did this while keeping fixed all the other parameters of the system. The choice of 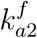 as one of the parameters to be scanned over for the bistability analysis is arbitrary and a similar parameter scan could be performed by choosing the other parameters as well and the results for bistability would still hold true. However, as we later show using sensitivity analysis, 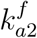 is also one of the sensitive parameters in the models, thereby showing a restrictive range for the bistability regime (as shown in Fig. 2).

The parameter scan confirmed the presence of bistability in our model. As mentioned earlier, bistablility leads to the threshold phenomenon where the system crosses the barrier and in this case goes to the higher aggregated state which could be lethal for the cell. In the parameter scan, shown in Fig. 2, the white region depicts the bistable region where the system has 3 steady state solutions, where, two states (lowest and highest values) represent the stable steady states and the one in the middle, the unstable steady state.

#### Stochastic Simulations

Once we had the bistable region identified from Fig. 2, we could choose parameters that would show either a single or a bimodal distribution of folded and aggregate proteins. As described in the previous section, we laid down our model to study the role of HSP, HSF and the HSP-HSF complex in the bistability that the system exhibits. In Model I, HSP and HSF are conserved in the system.

The first set of simulations, as shown in Fig. 3, were performed using total HSP counts = 2050, total proteins = 5000 and HSF = 1000, which are typical values [25]. The simulations resulted in a transient state for short time and eventually went to the higher aggregated state. The simulations were run for 1000 replicates in order to get a smooth distribution function. From Fig. 3(B), we could see that some replicates reach higher aggregated state sooner than others. From the probability distribution in Fig. 3(C), we could infer that the probability for the system to be in lower folded protein concentration is higher than that of higher folded protein concentration.

**FIG. 3:**
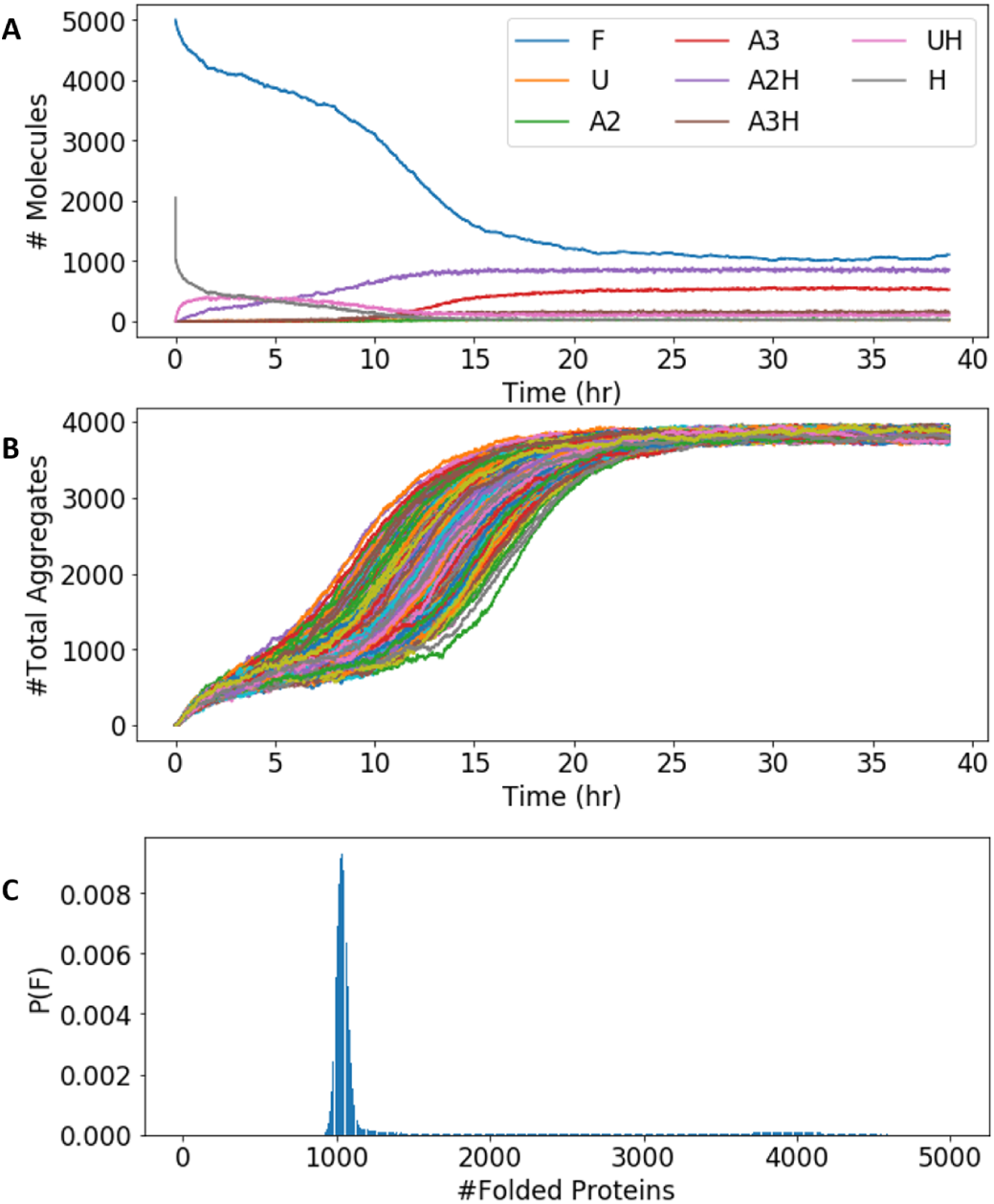
Simulations for 1000 replicates of Model I for conserved HSP and HSF. (A) The presence of bistability for the entities present in the system for one replicate. (B) Depiction of the stochasticity in the system and bistable behaviour for the formation of aggregates. (C) The Probability distribution function for 1000 replicates. Total Protein=5000, Total HSP=2050, Total HSF= 1000

Later, a second set of simulations was performed to see the role of HSP in the system. We hypothesized that the presence of more HSP would help the system to remain healthy and it would spend more time in folded state and after a definite time would cross the barrier and go to the higher aggregated state. Therefore, to test this hypothesis, we increased the HSP count to 2100 in our simulation of the system. This is shown in Fig 4. The simulations tested positive for our hypothesis. Indeed, increasing HSP in the system helped it to be in lower aggregated state. The system stayed in the lower aggregated state for a long duration of time, which is validated from the probability distribution curve where most of the proteins stayed in folded state.

**FIG. 4:**
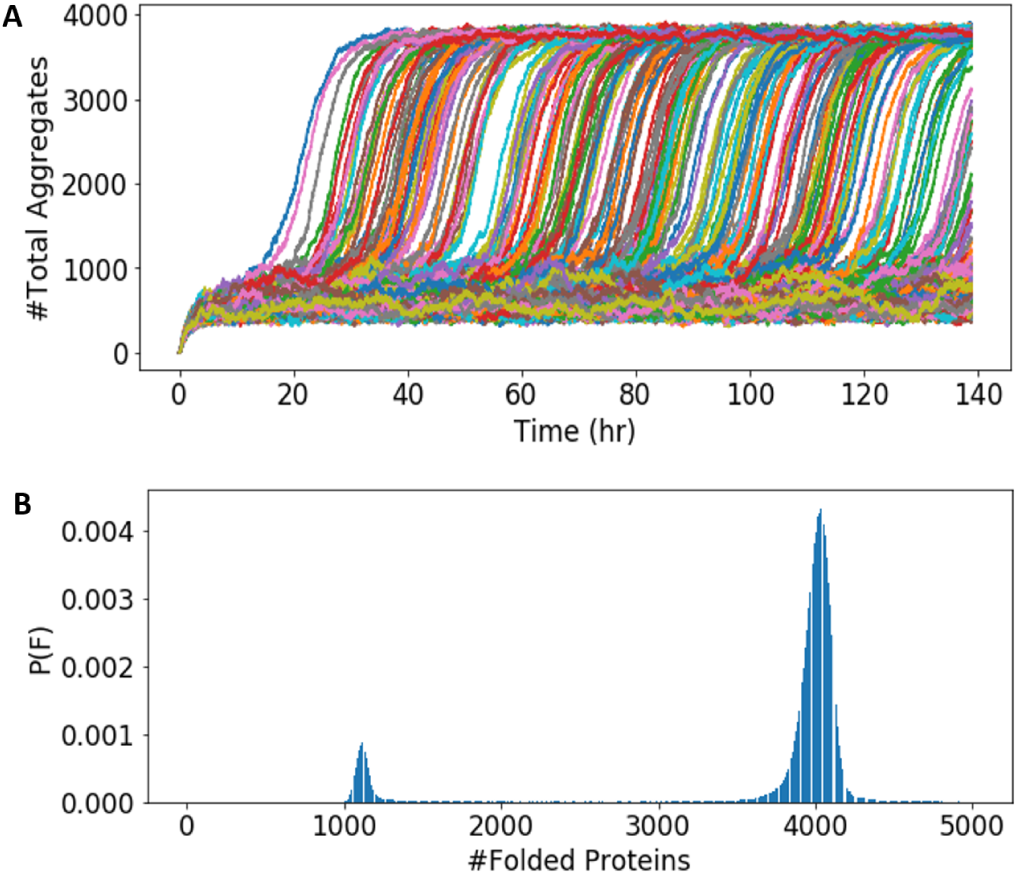
Simulations for 1000 replicates of Model I for conserved HSP and HSF. (A) Depiction of stochasticity in the system and the exhibition of bistability in the formation of aggregates. (B) The Probability distribution function of folded proteins for a collection of 1000 replicates. Total Protein=5000, Total HSP=2100, Total HSF= 1000

The simulations of Model I rightly represent the scenario that increasing the amount of HSPs increase the concentration of the folded protein compared to the unfolded protein, whereas, some of the proteins remain in the aggregated or unfolded state even at higher concentrations of HSP. However, in a real cell, the aggregated proteins do become folded under increasing concentrations of HSP. We addressed this drawback of Model I in Model II.

### B. Model II

The previous model was devised under the assumption that the HSPs are conserved. However, in the real system, the HSPs are produced continuously with increase in stress. This phenomenon was addressed in this model. Therefore, as listed in Table I, Model II has an additional reaction where HSP (H) is created dynamically from HSF (S). This means that the total HSP and the total HSF are no longer conserved which implies that Equations (3) and (4) are no longer valid for Model II. We built on the results of Model I and further explored this dynamical system through stochastic simulations.

#### Stochastic Simulations

As per the response mechanism, unfolded proteins tend to aggregate with an increase in the cell’s exposure to heat stress. The HSPs are chaperones that help fold back these unfolded proteins. The cells therefore, translate HSP as a response to their fight against such a stress. Fig. 5 shows the threshold phenomenon inside the cell under stress. Under stress, there is rapid misfolding and aggregate formation, but the presence of basal level HSP as an initial response of the cell tends to gradually decrease the aggregates and after a certain time due to the Heat Shock Response, the system crosses the threshold barrier and goes to the state of lower aggregates. The threshold barrier is crossed as the system reaches a certain level of HSP concentration. In this model, the HSPs are produced from the HSFs. In the real scenario, it is this HSF in its activated form that acts as transcription factors and translates HSP. The initial HSFs in the system are consumed to produce more HSP which helps in protein folding. Due to the gradual decrease in aggregate concentration, there is a wider distribution of folded proteins in the lower state i.e, around 1000 folded proteins (Fig. 5(C)).

**FIG. 5:**
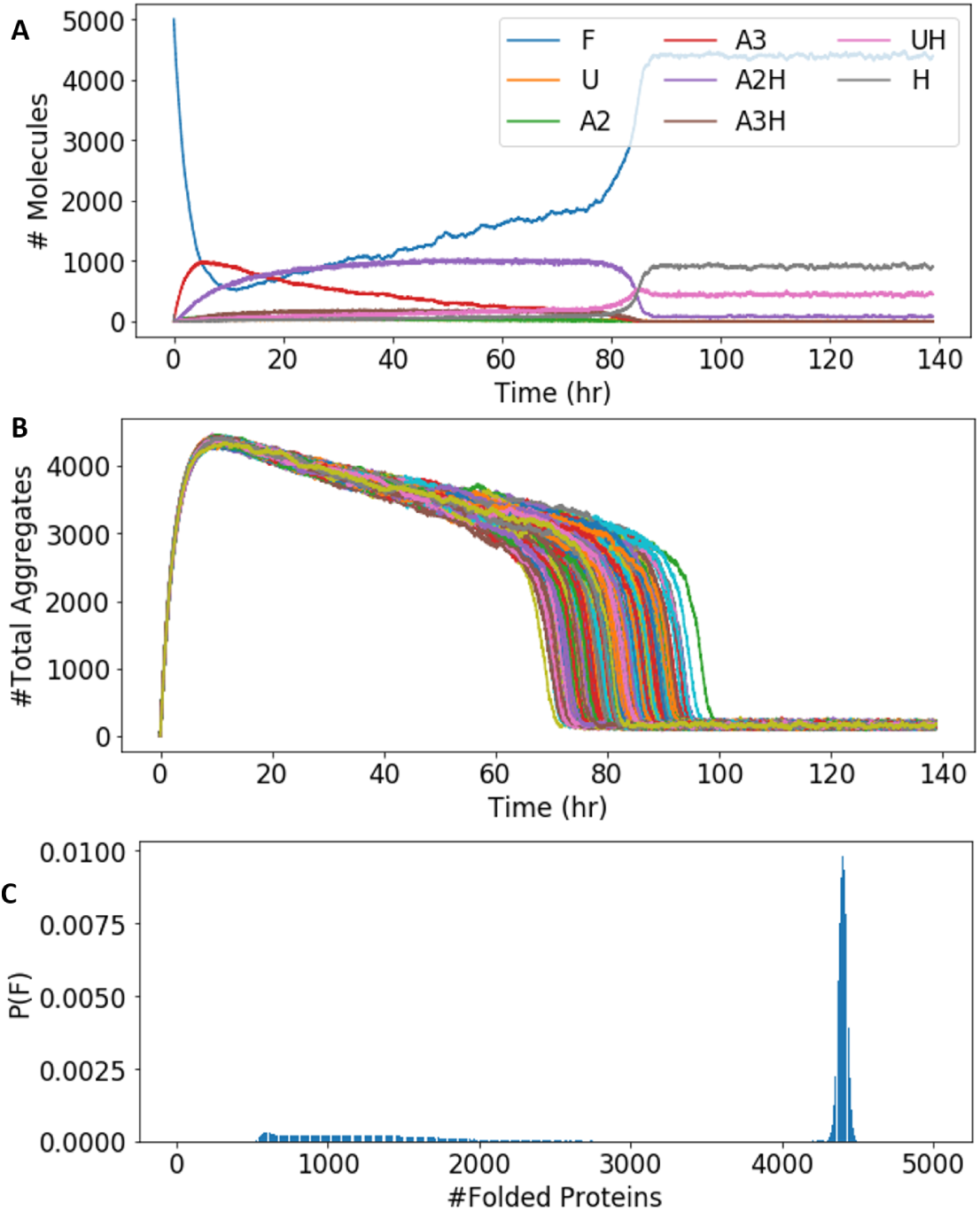
Simulations for 1000 replicates of the Heat Shock Response Model (Model II). (A)The presence of bistability for the entities present in the system for one replicate (B) Depiction of stochasticity in the system and the exhibition of bistability in the dissolution of aggregates. (C) The Probability distribution function for 1000 replicates. Total Protein=5000, Total initial basal HSP=100,Total initial HSF= 1500

In the simulations depicted in Fig. 5, we assumed that there are no HSP-HSF complex at the beginning of the simulation. But, the system could exist in the form where all the HSPs and HSFs are either segregated or in the form of a complex. Therefore, we next carried out simulations where the system in its initial state had all the HSPs in the form of the HSP-HSF complex. Since it had higher HSP concentration initially, the system immediately went to the lower aggregate state as shown in Fig 6(A). However, it is to be noted that eventually simulations depicted in both Figs. 5 and 6 had nearly equal amounts of HSP (counts 1000) in the final state. Further, Fig. 6(B) shows the probability distribution for the folded proteins in the system. The probability of finding approximately 4500 proteins in the folded state is higher, thereby depicting the healthy condition of the cell.

**FIG. 6:**
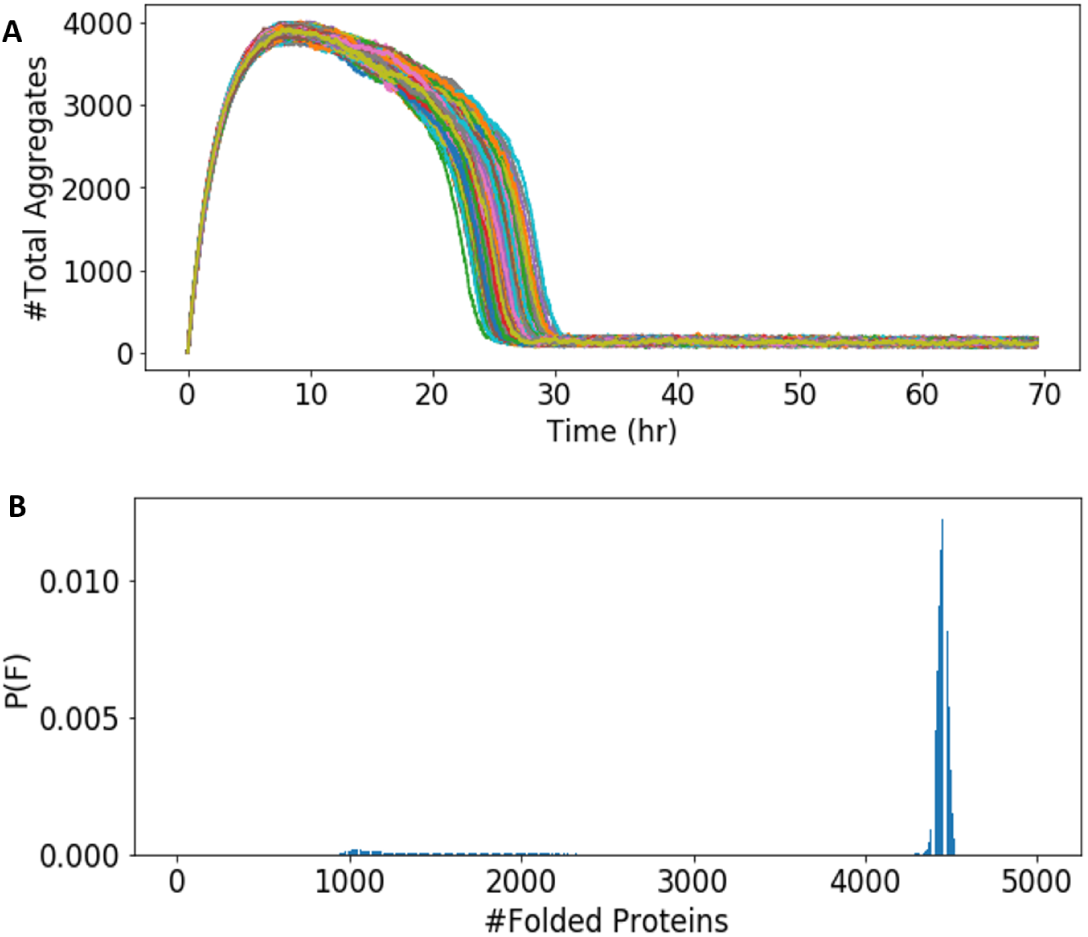
Simulations of Model II with HSF in the form of HSF-HSP complex initially. (A)1000 replicates for the system where all HSF and HSP are in the bound state.(B) Probability distribution curve for Total Protein count=5000, Free HSF=0, Initial HSF:HSP=1500, Initial Basal level HSP=100

We next studied the role of the basal level of HSP in the system. The amount of the basal level of HSP present decides the response time for the cell. To study this, we simulated our system with (HSP count=100) and without basal level HSP, shown in Figs. 7(A) and 7(B) respectively. It was evident that the cell tried to be in slowly decreasing aggregated steady state for longer when we had no HSP initially. This indicated a delayed response in this scenario.

**FIG. 7:**
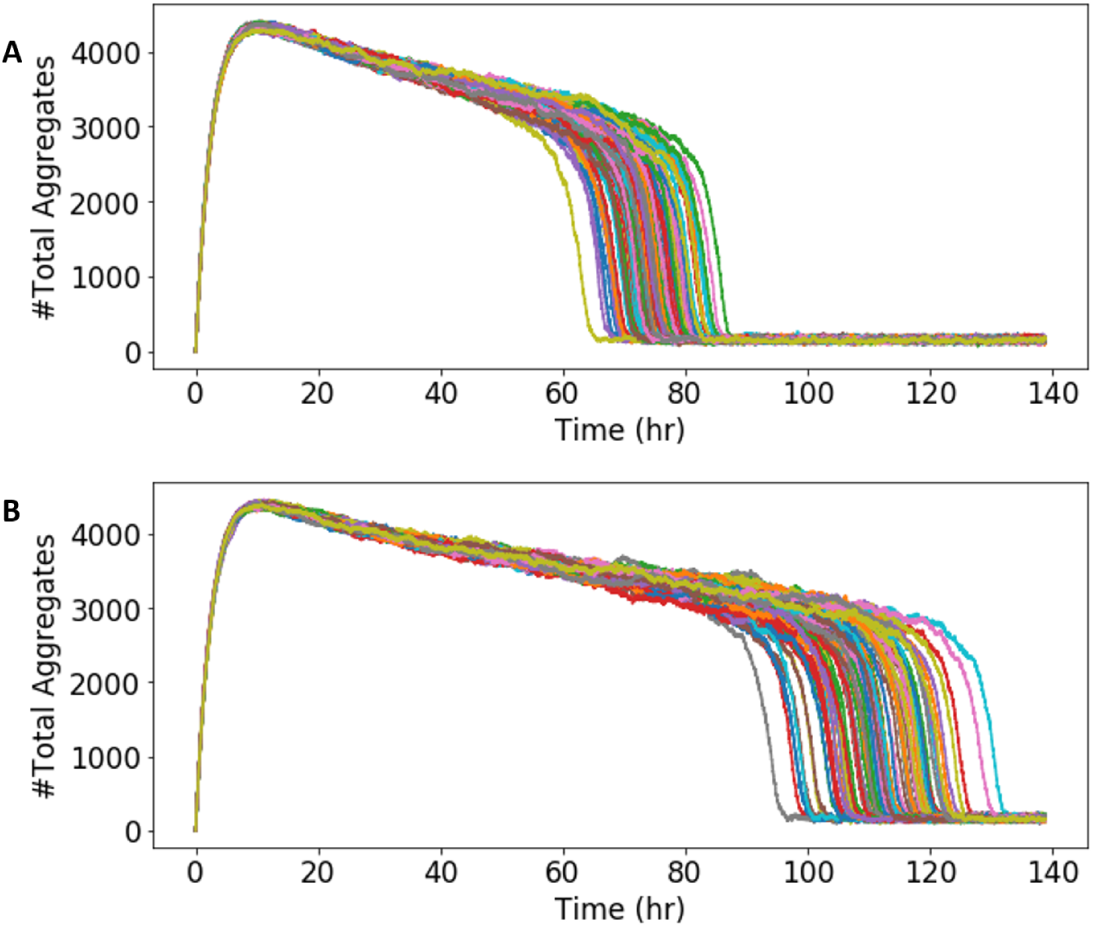
Significance of basal HSP. Simulations for 100 replicates (A) with basal HSP (counts = 100) and (B) without basal HSP (counts = 0) in the system. In both these simulations, total Protein=5000 and Initial HSF=1500.

### C. Sensitivity Analysis

To find the relationship between the species and the parameters used in our model, we estimated the sensitivity coefficients for every species and parameter [42]. Carrying out sensitivity analysis gives a quantitative estimate to questions such as which parameters are of utmost importance and which ones do not affect the model. It also gives a quantitative measure of how changes in different parameters of the model can potentially alter the processes inside the cell. Since we were only interested in studying the steady state kinetics of the bistable system, we calculated the sensitivity coefficient using Eq. 12.

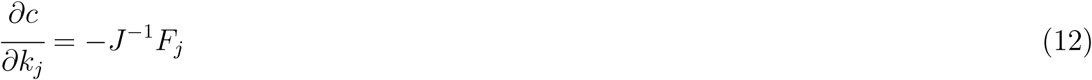

where *J* is the Jacobian Matrix and *F*_*j*_ is the *j*^*th*^ column of the matrix 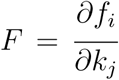. Here, *f*_*i*_ denotes the *i*^*th*^ function for the corresponding steady state differential equation with 1 ≤ *i* ≤ 8, c denotes the concentration of the species and *k*_*j*_ denotes the *j*^*th*^ parameter with 1 ≤ *j* ≤ 18.

The stationary sensitivity coefficient represents the change of stationary species concentration as a result of a differential change in parameters. After close introspection of all the sensitivity coefficients, the least sensitive parameter was 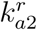 whose sensitivity coefficient value was way less than 0.1 for all the species in our system. To test this, we changed the value of 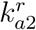 with an increment of 10% and tested the steady state values. Since, this was the least sensitive parameter, the values of none of the species changed significantly. The sensitivity coefficient for self refolding turned out to be negligible in our analysis. This further supported our assumption that the rate constants 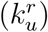 for self folding of mis-folded proteins were significantly smaller than the catalytic refolding. For e.g, when we compared the sensitivity coefficients for mis-folding and self refolding, we observed that the parameter 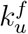 was more sensitive than 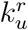. The 10 fold increase in the parameter 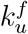 changed the protein counts of folded proteins from 1800 to 150 whereas, a 10 fold increase in 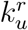 changed the protein counts from 1800 to 1900 only. Further, the parameter 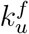 is temperature dependent (Eq. 1), thereby experiencing the direct consequence of heat stress.

The proper cellular functioning can only be performed by the cells that contain folded proteins. All the response that is mounted in the system drives the reactions towards folded state of the protein. Also, we assume that initially all the proteins are in their healthy state i.e, the folded state. Therefore, the folded proteins tend to be affected the most during the Heat Shock Response where almost all the parameters tend to affect its existence.

The parameter 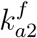 is a sensitive parameter which means that the aggregation process is indeed an important step in our model. We also performed our parameter scan taking 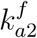 as one of the parameters to get our bistable regime. The other parameters which showed a similar trend in terms of sensitivity were 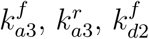 and 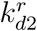.

In our simulations, we also studied the role of HSF and the HSF-HSP complex. The rate constants for these reactions directly affect the U:H complex and Folded proteins. Therefore, the reactions involving these components are significant. The kinetic rate constant, 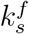, in fact, is a very sensitive parameter and corresponds to the production of the complex S:H which in turn produces UH and F. Therefore, any small increment in this parameter could lead to more folded proteins in the system. The S:H complex when bound to unfolded protein activates the response mechanism leading to a stable state consisting of higher concentration of folded proteins.

The complete set of sensitivity coefficients are listed in Table S1 in the supplementary information.

## DISCUSSION

We have, in this work, developed and carried out Monte Carlo simulations of two specific model systems that tie the HSR pathway with the various intermediate states of the protein folding and aggregation mechanism. In coming up with these models, we have tapped into two basic assumptions. One is that the total protein copy number is conserved due to the associated energy cost in increasing gene expression [44]. Also, proteins lost as irreversible aggregates due to very high stresses are replenished by translation which ensures protein homeostasis [24]. We have incorporated this phenomenon in our models and simulations in the form of Eq. 1. The second assumption is that chaperone assisted refolding kinetics is much faster than free refolding. This comes from many well studied systems and models which show that chaperones inhibit aggregation by binding to the exposed hydrophobic amino acid residues. Subsequently, these chaperone bound proteins go through a series of folding intermediates which have lowered energy barriers to get to the native state [14].

Following these features, we studied two specific models. Model I simply incorporated the HSF-HSP interaction and the related reactions while keeping the total HSP constant. Model II, on the other hand, in addition to the reactions of Model I also incorporated the fact that HSP is produced continuously with corresponding increase in stress. Model II is therefore closer to reality and this is reflected in the dynamical changes in the copy number of the folded and aggregated states of the protein. In the simulations of Model I (Figs. 3 and 4), due to reaction kinetic fluctuations, some of the proteins switched to the high aggregated state early. And since the total HSP was held constant, these high aggregated states once reached were long lived and highly stable. In Model II as well, stress did cause the folded proteins to become aggregated. But because of dynamical HSP concentration, as shown in the trajectories of Figs. 5A and 5C, all the proteins in the high aggregated state eventually re-folded. This also came about via the spontaneous transition characteristic of a bistable system, but as can be seen from Fig. 5C, there was a clear slowly decreasing aggregated state which was missing in Model I. Both, Models I and II, give a bimodal distribution as shown in Figs. 4B and 5C respectively. However, because of the slowly decreasing aggregated state, Model II gives a much broader distribution for the low folded (highly aggregated) state. The main advantage for the system to be explained by the latter model is that almost all the unfolded and aggregated proteins fold back to their native state. This is only possible because of the well-studied role of HSF in translating new HSPs [1].

The concentration of HSPs in the system is what determines the fate of the cell. Therefore, with a varying HSP concentration, we could observe a change in the response time of the cell. We define response time as the time at which the cell toggles between the two steady states in the bistable regime. For Model I, we calculated the average time for 1000 replicates to switch from one state to another. With HSP count=2050, it took about 10 hours to respond and switch into a higher aggregated state whereas, an increase in the HSP count by 50, doubled the response time for folded to aggregated state. Hence, these response times were consistent to our theory i.e, the HSP concentration affects the amount of time the cell stays in one particular state.

Finally, as shown in Figs. 6 and 7, we also extended the features of Model II to study the role of free versus HSP bound HSF and the role of basal HSP respectively. It is clear that initial high concentration of HSP either in the form of the HSF-HSP complex (Fig. 6) or in the form of free basal HSP (Fig. 7) leads to faster refolding dynamics. This is because the aggregated proteins then do not have to wait as long for the synthesis of new HSPs. Therefore, the re-folding process can start earlier. However, as can be seen, both from Figs. 6 and 7, there is some time lag in the response to be established and the aggregated state of the proteins are stable for a long time. The response time can be reduced by changing the number of HSFs, HSPs and the kinetic rate for the production of HSP from HSF. This could be observed in the simulation studies shown in Fig. 8. Initially, as per the parameter range listed in Table II, we changed the rate constant, *k*_*m*_, for the production of HSPs. Since, *k*_*m*_ is an important parameter for the cell to mount a response, its value [28, 35, 38] plays a key role in setting the response time. The wide range of *k*_*m*_ could be a reflection of the different forms of HSPs that are part of the HSR mechanism. We therefore increased the value of *k*_*m*_ by an order of magnitude and observed a significant change in the response time between the simulation trajectories shown in Figs. 8(A) and 8(B). On further increasing the value of *k*_*m*_ from 0.001*s*^−1^ to 0.005*s*^−1^, we could clearly see a decrease in the response time by several hours, as shown in Figs. 8(C) and 8(D), even though the number of HFs (S) in the system is reduced to 1400. As mentioned before, the state of the cell can also change depending on the nature of molecules present inside it. For example, HSPs and HSFs could either be present in the free form or as a complex. We simulated a case where all the HSFs are in the form of the HSF:HSP complex. By changing the amount of HSF:HSP complex, there is a change in the HSP molecules which affects the response time inside the cell. The presence of 900 (1500 to 2400) more HSPs inside the cell, even at a low *k*_*m*_ = 0.0001, reduced the response time by several hours. This comparison is shown in Figs. 8(E) and 8(F). Such an effect of increased HSF that we predict here can be tested experimentally by overexpressing HSF in the cell. Further, this kind of time lag between stress and response is a general phenomenon observed in several model organisms. For example, in yeast, delayed proteomic response is observed under severe heat stress leading to delayed HSR [24]. Similar delayed response was also observed in *C. elegans* under 34 deg Celsius high heat stress for 1 hour [3].

**FIG. 8:**
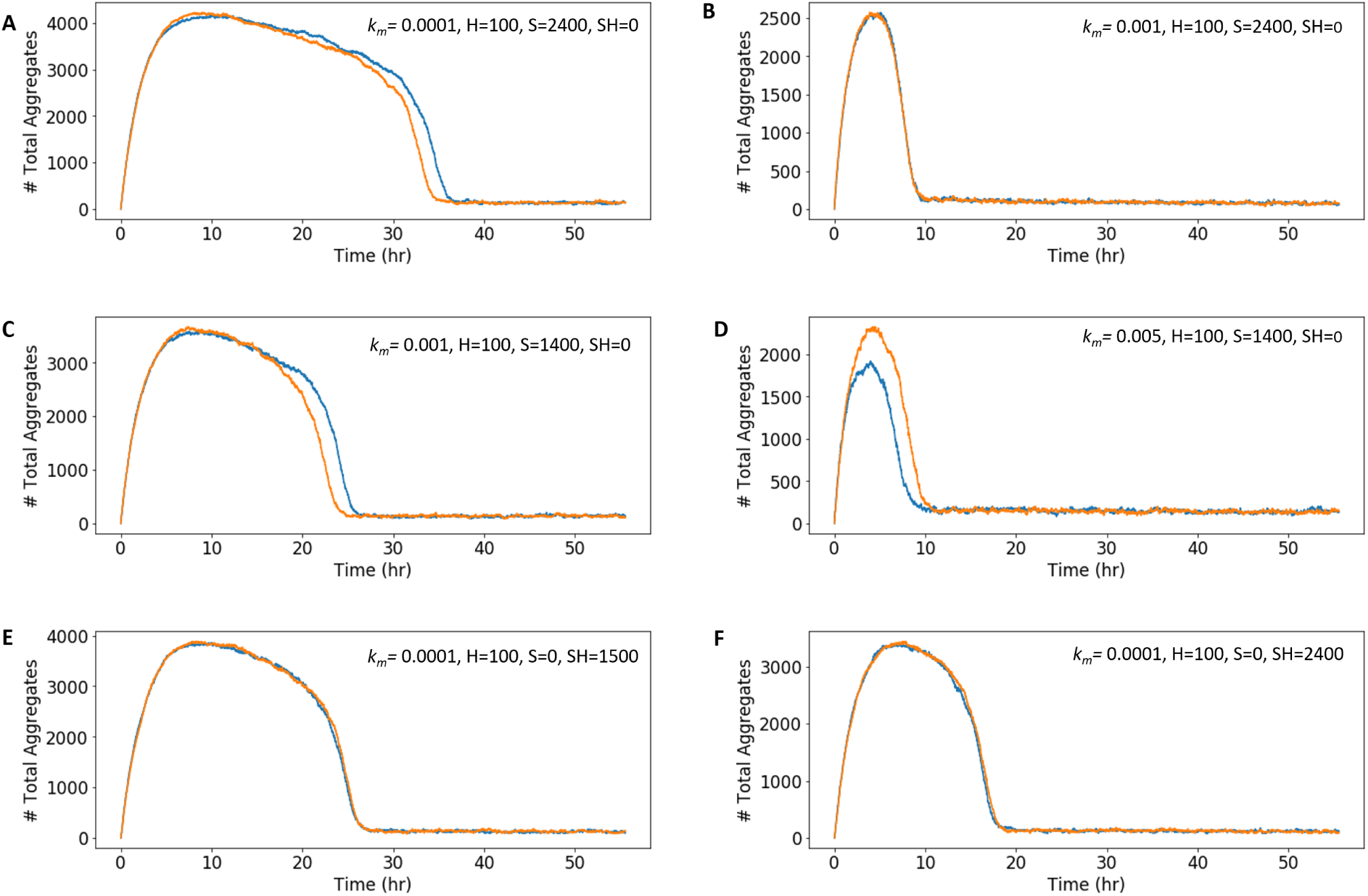
Role of HSF in Model II. Each figure depicts the role of molecular counts of HSPs and HSFs and the rate of production of HSP on the response time. The stochastic simulations were carried out for 2 replicates. The value of *k*_*m*_ is in the units *s*^−1^. We compare Fig. (A) with (B), Fig. (C) with (D) and Fig. (E) with (F).

In conclusion, the study of the two models revealed novel features regarding the specific roles of HSF, the HSF-HSP complex and the synthesis of HSP under stress in tipping the balance between folded and aggregated states. This would not have been possible without connecting the HSR mechanism to the protein aggregation framework. Therefore, future exploration of stress response mechanisms should be directed towards connecting related pathways. This might make the system more complicated, adding several more reactions and parameters, but will eventually give better insights into the larger inter-connectivity of signaling pathways.

## ACKNOWLEDGMENTS

Sushmita Pal is supported by the National Fellowship INSPIRE (Innovation in Science Pursuit of Inspired Research) funded by the Department of Science and Technology (DST), Government of India. This work has also been funded by the Science and Engineering Research Board (SERB) Startup Research Grant (Ref. No. SRG/2019/000726) awarded by DST, India.

