## Supplementary material for "A mathematical model of heat shock response to study the competition between protein folding and aggregation"

### Supplementary Information

| Sensitivity Coefficients | Value | Sensitivity Coefficients | Value |
|---|---|---|---|
| $\frac{\partial \ln f}{\partial \ln k_u^f}$ | -1 | $\frac{\partial \ln a_2 h}{\partial \ln k_{a_3}^r}$ | 0.530 |
| $\frac{\partial \ln f}{\partial \ln k_{a_2}^f}$ | -0.189 | $\frac{\partial \ln f}{\partial \ln k_{d_3}^f}$ | -0.478 |
| $\frac{\partial \ln u}{\partial \ln k_{a_2}^f}$ | -0.221 | $\frac{\partial \ln u}{\partial \ln k_{d_3}^f}$ | -0.485 |
| $\frac{\partial \ln a_2}{\partial \ln k_{a_2}^f}$ | 0.319 | $\frac{\partial \ln a_2}{\partial \ln k_{d_3}^f}$ | -0.557 |
| $\frac{\partial \ln uh}{\partial \ln k_{a_2}^f}$ | -0.189 | $\frac{\partial \ln a_3}{\partial \ln k_{d_3}^f}$ | -0.988 |
| $\frac{\partial \ln a_2 h}{\partial \ln k_{a_2}^f}$ | 0.319 | $\frac{\partial \ln uh}{\partial \ln k_{d_3}^f}$ | -0.478 |
| $\frac{\partial \ln f}{\partial \ln k_{a_3}^f}$ | -0.550 | $\frac{\partial \ln a_2 h}{\partial \ln k_{d_3}^f}$ | -0.556 |
| $\frac{\partial \ln u}{\partial \ln k_{a_3}^f}$ | -0.557 | $\frac{\partial \ln f}{\partial \ln k_{a_3}^r}$ | -0.214 |
| $\frac{\partial \ln a_2}{\partial \ln k_{a_3}^f}$ | -0.638 | $\frac{\partial \ln u}{\partial \ln k_{d_3}^r}$ | -0.207 |
| $\frac{\partial \ln uh}{\partial \ln k_{a_3}^f}$ | -0.550 | $\frac{\partial \ln uh}{\partial \ln k_{d_3}^r}$ | -0.214 |
| $\frac{\partial \ln a_2 h}{\partial \ln k_{a_3}^f}$ | -0.638 | $\frac{\partial \ln a_3 h}{\partial \ln k_{d_3}^r}$ | -0.973 |
| $\frac{\partial \ln f}{\partial \ln k_{a_3}^r}$ | 0.485 | $\frac{\partial \ln f}{\partial \ln k_{d_2}^f}$ | 0.506 |
| $\frac{\partial \ln u}{\partial \ln k_{a_3}^r}$ | 0.492 | $\frac{\partial \ln u}{\partial \ln k_{d_2}^f}$ | 0.506 |
| $\frac{\partial \ln a_2}{\partial \ln k_{a_3}^r}$ | 0.563 | $\frac{\partial \ln a_2}{\partial \ln k_{d_2}^f}$ | -0.287 |
| $\frac{\partial \ln uh}{\partial \ln k_{a_3}^r}$ | 0.485 | $\frac{\partial \ln uh}{\partial \ln k_{d_2}^f}$ | 0.506 |

| Sensitivity Coefficients | Value | Sensitivity Coefficients | Value |
|---|---|---|---|
| $\frac{\partial \ln a_2 h}{\partial \ln k^f_{d_2}}$ | 0.711 | $\frac{\partial \ln uh}{\partial \ln k_{d_2}}$ | 0.491 |
| $\frac{\partial \ln f}{\partial \ln k^r_{d_2}}$ | -0.491 | $\frac{\partial \ln a_2 h}{\partial \ln k_{d_2}}$ | -0.308 |
| $\frac{\partial \ln u}{\partial \ln k^r_{d_2}}$ | -0.491 | $\frac{\partial \ln f}{\partial \ln k_f}$ | 0.988 |
| $\frac{\partial \ln a_2}{\partial \ln k^r_{d_2}}$ | 0.279 | $\frac{\partial \ln f}{\partial \ln k^f_s}$ | 1.18 |
| $\frac{\partial \ln uh}{\partial \ln k^r_{d_2}}$ | -0.491 | $\frac{\partial \ln uh}{\partial \ln k^f_s}$ | 1.18 |
| $\frac{\partial \ln a_2 h}{\partial \ln k^r_{d_2}}$ | -0.691 | $\frac{\partial \ln sh}{\partial \ln k^f_s}$ | 7.46 |
| $\frac{\partial \ln f}{\partial \ln k^f_f}$ | 0.768 | $\frac{\partial \ln f}{\partial \ln k^r_s}$ | -0.158 |
| $\frac{\partial \ln uh}{\partial \ln k^f_f}$ | 0.769 | $\frac{\partial \ln uh}{\partial \ln k^r_s}$ | -0.158 |
| $\frac{\partial \ln f}{\partial \ln k^r_f}$ | -0.988 | $\frac{\partial \ln sh}{\partial \ln k^r_s}$ | 0.999 |
| $\frac{\partial \ln uh}{\partial \ln k^r_f}$ | -0.990 | $\frac{\partial \ln f}{\partial \ln k^u_s}$ | 0.158 |
| $\frac{\partial \ln f}{\partial \ln k_{d_3}}$ | 0.207 | $\frac{\partial \ln uh}{\partial \ln k^u_s}$ | 0.158 |
| $\frac{\partial \ln u}{\partial \ln k_{d_3}}$ | 0.200 | $\frac{\partial \ln a_2}{\partial \ln k_{d_2}}$ | -0.279 |
| $\frac{\partial \ln uh}{\partial \ln k_{d_3}}$ | 0.207 | $\frac{\partial \ln u}{\partial \ln k_{d_2}}$ | 0.491 |
| $\frac{\partial \ln f}{\partial \ln k_{d_2}}$ | 0.491 | | |

Table 1: Sensitivity Coefficients
